# Studying sleep orthologs in Epsilonproteobacteria through an evolutionary lens: Investigating sleep mysteries through phylogenomics

**DOI:** 10.1101/2025.01.02.630604

**Authors:** Seithikurippu R Pandi-Perumal, Konda Mani Saravanan, Sayan Paul, David Warren Spence, Saravana Babu Chidambaram

**Affiliations:** Department of Pharmacology, JSS College of Pharmacy, JSS Academy of Higher Education & Research, Mysuru 570015, Karnataka, India; Centre for Research and Development, Chandigarh University, Mohali, Punjab, 140413, India; Division of Research and Development, Lovely Professional University, Phagwara, Punjab, 144411, India; Department of Biotechnology, Bharath Institute of Higher Education and Research, Chennai, Tamil Nadu, 600073, India; Department of Biochemistry & Molecular Biology, The University of Texas Medical Branch at Galveston, Galveston, TX, 77555, USA; Research Scientist, Dufferin Street, Toronto, ON M6K 2B4, Canada; Department of Pharmacology, JSS College of Pharmacy, JSS Academy of Higher Education and Research, Mysuru, Karnataka, 570015, India; Special Interest Group - Brain, Behaviour and Cognitive Neurosciences, JSS Academy of Higher Education & Research, Mysuru, Karnataka, 570015, India

**Keywords:** Epsilonproteobacteria, Orthologous Genes, Phylogenomics, Potassium Channels, Sleep, *Sulfurimonas Paralvinellae*

## Abstract

The current study employed phylogenomic methods to examine the evolutionary role and significance of sleep-related genes in *Sulfurimonas paralvinellae* of the Epsilonproteobacteria class. This has facilitated the identification of conserved sleep orthologs, including DnaK, serine hydroxymethyltransferase, and potassium channel family proteins, exhibiting sequence similarities ranging from 39.13% to 61.45%. These findings align with prior research indicating that chaperones and ion channels are conserved during sleep. This was demonstrated by the observation that proteins with fewer domains exhibited more significant conservation than others, such as adenylate kinase, which is substantial under selective pressure. Distinct adaptations in bifunctional protein - serine/threonine kinases and phosphatases were linked to *S. paralvinellae*, an extremophilic organism adapted to high-pressure and/or high-temperature conditions, indicating functional divergence influenced by the organism’s environment. The Gene Ontology study results indicated catalytic activity, potassium channel function, and cellular processes, underscoring the significance of ion channels in regulating the sleep-wake cycle. Furthermore, the categories not recognized as particularly significant for the over-represented genes encompassed metabolic and signal transduction categories, suggesting enhanced functional flexibility within this protein subfamily. The findings emphasize that orthologous interactions are complex and influenced by subfunctionalization and neofunctionalization of ecology and evolution. These findings enhance the existing understanding of the evolution of sleep-related genes and their association with metabolic and environmental changes, providing a foundation for subsequent experimental investigations and cross-taxonomic comparisons.

## INTRODUCTION

The field of sleep science has grown significantly in recent years, and with it, the nature of both normal and pathological sleep processes have been increasingly characterized. For instance, The International Classification of Sleep Disorders (ICSD), Third Edition, Text Revision, now recognizes over a hundred different sleep disorders. Despite its phenomenal growth in recent years, the field of sleep science has nevertheless tended to focus on the behavioral and neurophysiological aspects of sleep while much of its genetic basis remains poorly understood. Sleep is a complex and highly conserved behavior that has been viewed as an evolutionary conundrum, inasmuch as the near universality of its occurrence suggests that sleep is important for survival, but its exact function remains a biological mystery [1]. Most animals that have been sufficiently investigated to this point exhibit sleep or behaviors resembling sleep, and, further, sleep appears to play a crucial role in their daily routine. Although sleep can be attributed to a wide range of physiological processes, the evolutionary purpose of sleep is still unclear, at least at the genetic and molecular levels [2].

The evolutionary origins and function of sleep are among the two greatest mysteries in biology. In this regard, hundreds of species from different taxa have had their sleep characteristics studied, although firm conclusions regarding the meaning of these characteristics have not been drawn [3–6]. Moreover, many of the studies are difficult to compare due to differences in their respective scope, depth, and focus.

Most early research into basic sleep mechanisms began by examining sleep in animal models, including rodents, cats, dogs, and non-human primates. Over the last few decades, the effective use of simple animal models to explain sleep has emerged [7]. Among the examples of model organisms that have been extensively utilized in sleep research have been *Drosophila melanogaster* (fruit fly), *Caenorhabditis elegans* (a transparent nematode), and *Danio rerio* (Zebrafish) [8]. Additionally, sleep has been documented in other primitive organisms such as the brainless freshwater polyp *Hydra vulgaris* (cnidarian) [9] and Jellyfish Cassiopea, which exhibits a ‘sleep-like’ state [10].

Bergey’s Manual of Systematic Bacteriology (Second Edition, Volume 2, Part C: Alpha-, Beta-, Delta-, and Epsilonproteobacteria) states that the classification of the class Alphaproteobacteria was determined using 16S (known as the small subunit) Ribosomal ribonucleic acid (rRNA) gene sequences, which has emerged as a gold standard technique in phylogenetic taxonomic analysis. *Campylobacteraceae*, *Helicobacteraceae*, and *Nautilaceae* are the families that make up the order Campylobacterales, composed of the class Epsilonproteobacteria.

Epsilonproteobacteria is a widely distributed class of flagellated bacteria commonly linked to hydrothermal vents or the digestive tracts of animals [11]. These organisms are often spiral or curved in shape and exhibit diverse metabolism and ecological niches. For example, Nautilia and Caminibacter are naturally anaerobic marine thermophiles.

It has been suggested that deep-sea hydrothermal vents, which are distinct habitats supporting extremely productive ecosystems powered by geochemical energy, may be the location of life’s beginning and early evolution [12]. For instance, a pH of 5.5 and vent fluid temperatures of 310–320 °C have been reported within the Loki’s Castle hydrothermal vent field on the Arctic Mid-Ocean Ridge (AMOR), about 73 degrees north off the coast of Greenland. The two most common bacterial groups found in the area were Aquificales and Epsilonproteobacteria members. We postulated that since alphaproteobacteria have been linked to the origin of mitochondria, Epsilonproteobacteria would be a promising group to investigate using phylogenomics. The phylogenomics approach, which was introduced by the present authors to analyse sleep genes, is more effective than the phylogenetic methodology. They enable one to get a much closer look at genetic phenomena in simple organisms.

Epsilonproteobacteria often show slender spiral rods with polar flagella. Another characteristic of this organism is that it can swim rapidly and continuously, even in harsh marine environments with high viscosity [11]. The current study investigated *Sulfurimonas Paralvinellae.* This epsilonproteobacteria possesses a genome comprising approximately 2.4 million base pairs [Genbank accession number, GCA_014905135.1]. This compact genome includes genes for sulfur oxidation and nitrate reduction, making it well-suited for the extreme conditions that often surround hydrothermal vents. The genes associated with this organism are linked to chemolithoautotrophic metabolism, stress responses, and various energy-generating pathways [13]. The genome also contains conserved molecular chaperones, ion transporters, and multifunctional enzymes such as bifunctional protein-serine/threonine kinases, indicating significant evolution and metabolic processes.

## METHODS

### Collection of sleep genes

Sleep genes were compiled using the NCBI database by utilizing search terms that included ‘gene,’ ‘sleep,’ ‘circadian rhythm,’ ‘clock,’ ‘insomnia,’ and ‘sleep regulation’ [14]. The filters, including “homo sapiens,” along with relevant years and research types, were applied to ensure the inclusion of only human genes. A literature survey was conducted to assess the direct involvement of each gene in sleep processes or disorders, utilizing functional annotations and scholarly articles from PubMed. The data were validated using GeneCards and Online Mendelian Inheritance in Man (OMIM) databases, and new information obtained was incorporated [15,16]. The final output is a comprehensive catalog of genes, including their symbols, full names, and annotations. This resource will be highly advantageous for any future research into the molecular etiology of sleep and sleep disorders. A similar procedure was utilized in our recent publication [17].

### Blast Analysis of Sleep Genes against *Sulfurimonas Paralvinellae* Genome

A BLAST analysis, accompanied by comparison research, was initially performed referencing the *Sulfurimonas paralvinellae* genome, which has sleep-related genes. Nucleotide or protein sequences of these genes were obtained from the NCBI database in FASTA format, utilizing the BLAST tool, with options for BLASTn for nucleotide sequences or BLASTp for protein sequences [18]. The target database was designated as the *Sulfurimonas paralvinellae* genome, which is available for download from NCBI or alternative sources. The sequences for comparison or analysis were either uploaded or entered directly into the “align two or more sequences” option. Parameters in the search were optimized, including E-value thresholds, match/mismatch scores, gap penalties, and word size. The importance of the alignments was assessed using scores, E-values, and identity percentages. These were used to identify matches and to judge the biological relevance of the similarities. The findings were further evaluated based on recent findings in the scientific literature. The aggregated data were then used to infer the functions of the orthologous genes in *S. paralvinellae*.

### Orthology and functional annotation of sleep genes

The public database eggNOG (http://eggnog6.embl.de/) contains functional annotations, gene evolutionary histories, and orthology linkages. The functional annotation (EggNOG) of sleep-related genes was performed using nucleotide or protein sequences obtained in FASTA format from the NCBI database and then examined using the EggNOG database [19]. The EggNOG-mapper application on the EggNOG website facilitated the analysis. The gene sequences were subsequently input into the EggNOG-mapper portal, utilizing specific search criteria by selecting the appropriate taxonomic range and adjusting parameters such as the E-value for the search. The computer overlaid sequences onto EggNOG orthologs, incorporating functional annotations based on conserved domains and established functions. The obtained data were analyzed for Gene Ontology (GO) keywords, as these appear in the Kyoto Encyclopedia of Genes and Genomes (KEGG) pathways [20], and Clusters of Orthologous Groups (COG) categories [21], with a primary emphasis on domains and pathways associated with sleep. The EggNOG findings were subsequently corroborated using a literature review, wherein the results were compared with prior investigations to authenticate the derived functional annotations. This comparative method permitted inferences into the biological functions and processes associated with sleep genes.

## INTERPRO SCAN OF SLEEP GENES

InterProScan analysis was conducted on sleep-related genes using sequences obtained from the NCBI database in FASTA format. The InterProScan tool (https://www.ebi.ac.uk/interpro/interproscan.html) predicts protein domains, families, and functional sites based on submitted sequences [22]. The search parameters were optimized to incorporate Pfam, PRINTS, and PROSITE patterns with user-defined settings. It integrated predictive models from various sources to discover conserved domains and active and protein families. The results were thoroughly analyzed to provide descriptions, with a comparative assessment with existing studies to elucidate the role of sleep genes in biological processes.

### Pathway analysis of sleep genes

The Gene ID or Gene name / Entrez ID of sleep-related genes was acquired for Pathway Analysis utilizing KEGG (https://www.genome.jp/kegg/), Reactome, IPA, or DAVID [23]. The gene list was rendered compatible, and the analysis parameters were established by selecting the organism “*Sulfurimonas paralvinellae*” and opting for pathway enrichment and network analysis. The research correlated the genes with established biological functions and signaling networks while identifying statistically overrepresented pathways. To evaluate the influence of the study results, evidence from the relevant genes, statistical characteristics, and the biological relevance of the pathways related to sleep mechanisms were contrasted with findings from previous research. The final report encompassed fundamental details, including input genes, analytical parameters, a compilation of enriched pathways, p-values, and graphical representations, including route diagrams and networks.

## RESULTS

### Sleep Genes BLAST Results Against *Sulfurimonas paralvinellae* Genome

Table 1 summarizes the protein BLAST search conducted to identify homologs of sleep-related genes in the genome of *S. paralvinellae*. Additional characteristics such as sequence length, e-value, and similarity mean were also evaluated. The e-value, indicating the degree of statistical significance [24], shows extremely significant hits; with e-values is 0, the proteins include molecular chaperone DnaK, succinate-CoA ligase subunit alpha, and translational GTPase TypA. The findings indicate significant similarities, suggesting that the proteins examined may be involved in processes analogous to sleep-related mechanisms seen in other animals. DnaK participates in processes such as protein folding and stress response, as well as cellular repair during sleep [25].

**Table 1.**
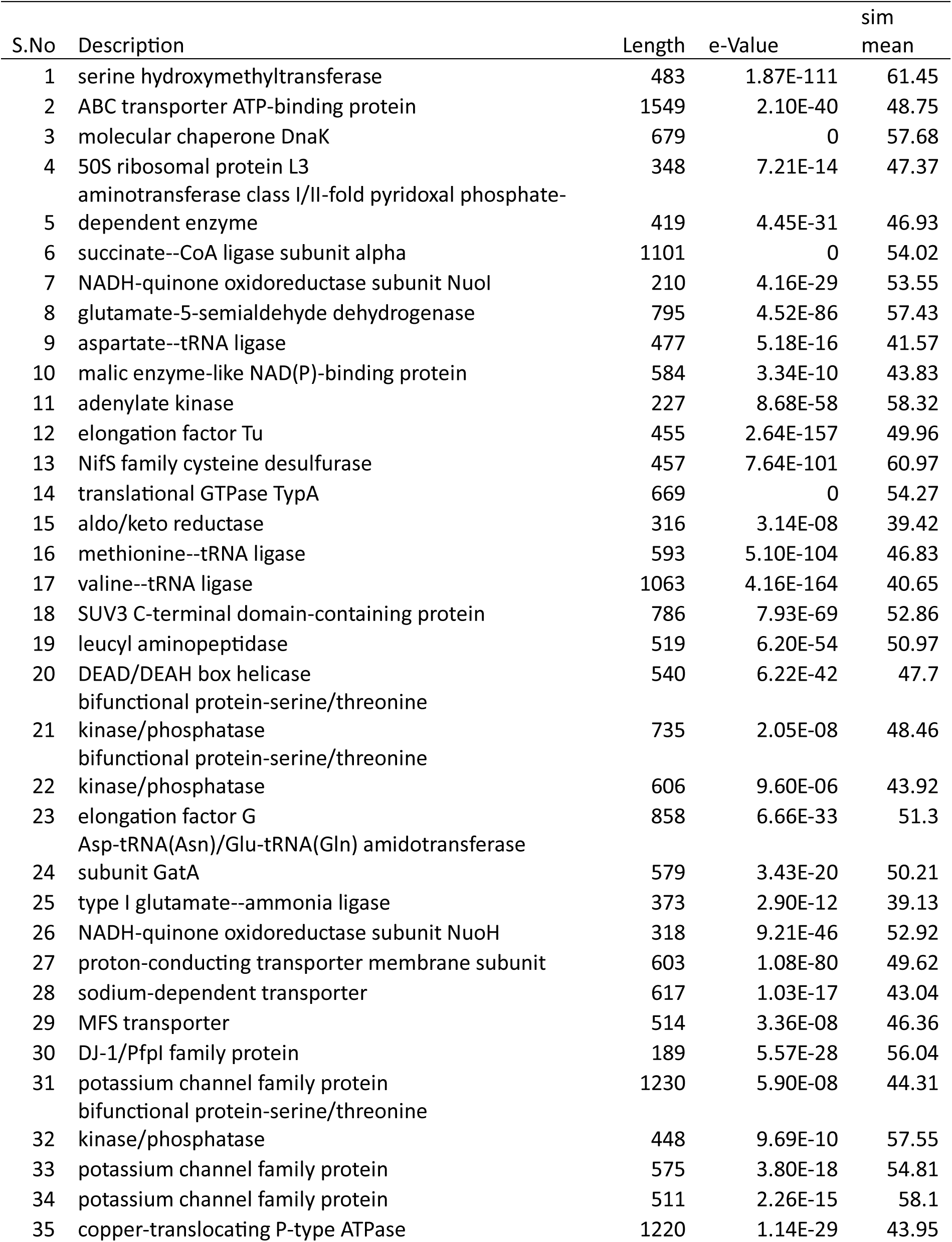
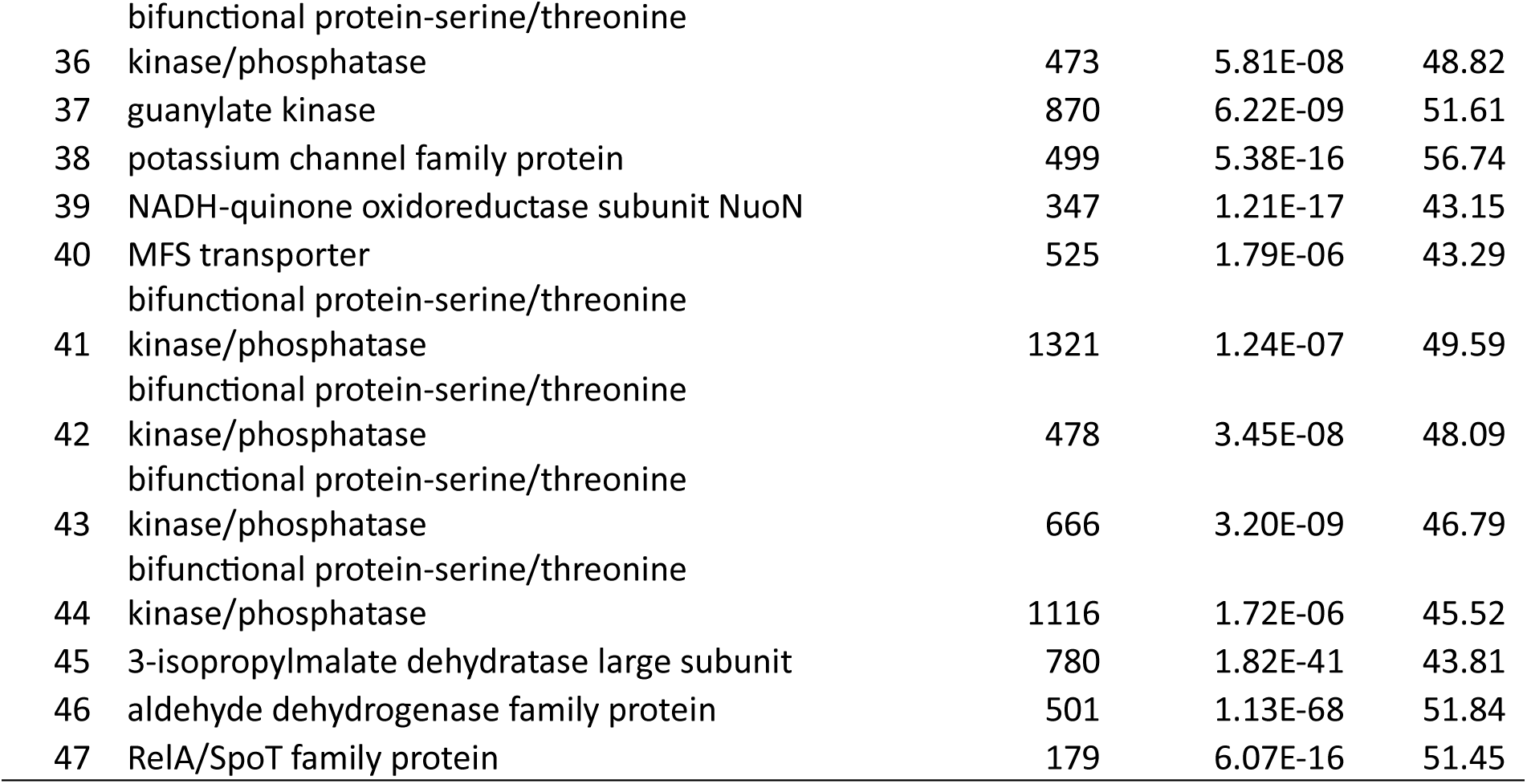
Hits obtained by performing sleep genes BLAST against *Sulfurimonas paralvinellae* genome.

All proteins possess identical sequence lengths and exhibit considerable similarity. The proportion of sequence similarity tends to be lower in longer proteins: the ABC transporter ATP-binding protein has 1549 amino acids, whereas the potassium channel family contains up to 1230 amino acids. This pattern indicates the inherent nature of multi-domain systems and seems rational. In contrast, shorter proteins like adenylate kinase (227 amino acids) have comparable mean similarity values of 58.32%, suggesting that their functional domains possess similar sequences across other species. The mean similarity values of these sets, ranging from 39.13 to 61.45, indicate moderate to high conservation of these proteins throughout the genome of *S. paralvinellae*, thus indicating their potential functional relevance.

A significant highlight is the recurrent identification of possible bifunctional protein-serine/threonine kinases and phosphatases [26]. Table 1 shows that these enzymes many times with varying sequence lengths and moderate similarity values (43.92–57.55). These proteins serve various functions and are involved in regulatory processes or signal transduction, which may be linked to environmental adaptation or adjustment of metabolic rates. These capabilities may mimic sleep-like states, suggesting their adaptive role in response to the particular environment of *S. paralvinellae*.

Figure 1 (top panel) presents the distribution of BLAST hits for the specified sleep-related genes and their sequences. The y-axis represents one variable, while the x-axis denotes the other, illustrating the hits and sequences to depict the patterns in the alignment results. The distribution likely indicates sequences corresponding with increased frequency, implying that the sleep genes in question are conserved or of functional significance. Differences in distribution may arise from variations in gene conservation or the availability of homologous sequences across different organisms.

**Figure 1.**
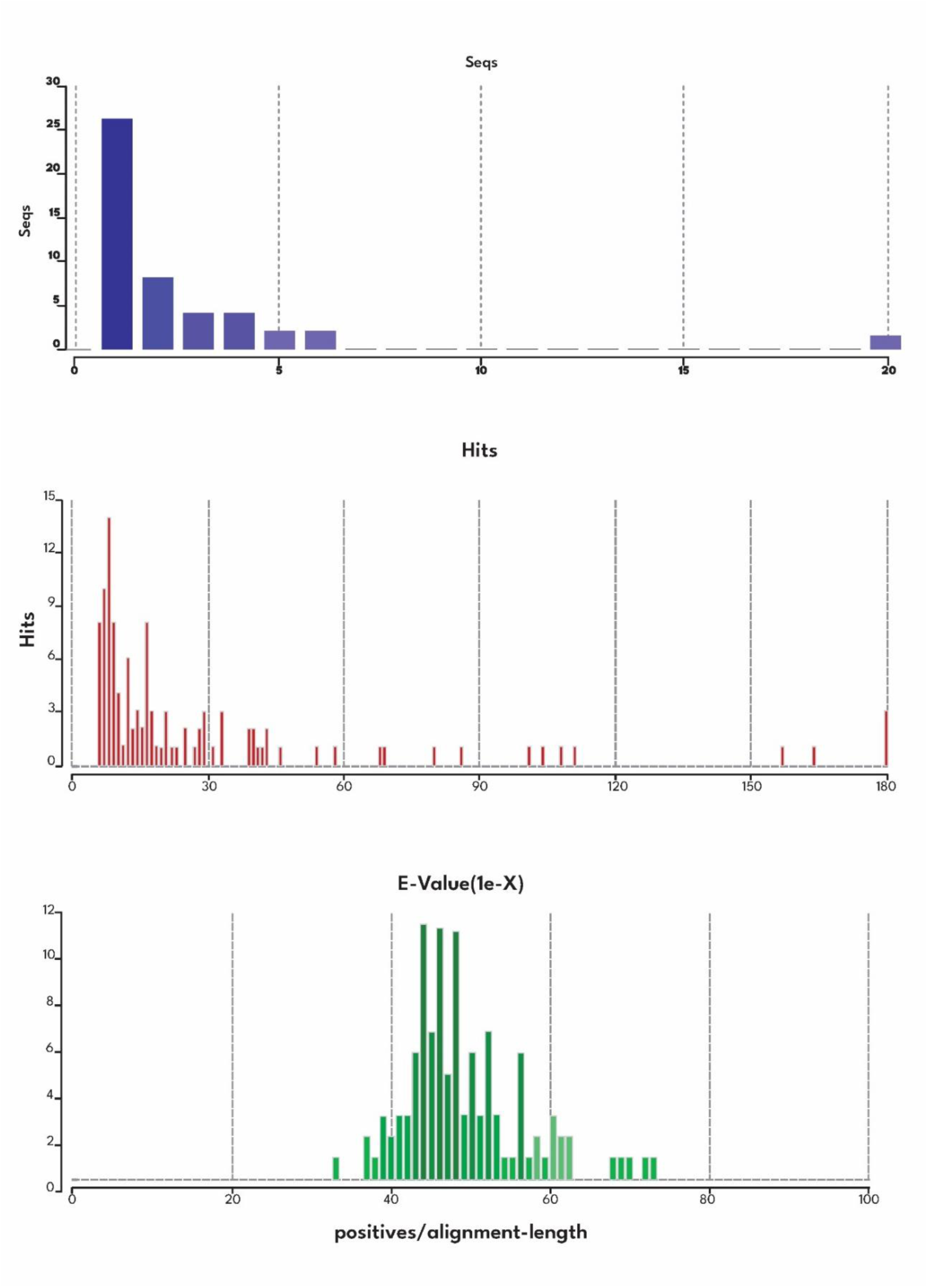
Analysis of sequence alignment statistics. The top panel displays the distribution of sequences (Seqs) across various bins, highlighting a high frequency of sequences within the lower range, with a notable peak at the initial bins and a smaller spike near the higher range. The middle panel shows the distribution of hits across alignment scores, demonstrating a concentration of hits in the lower ranges and a sparse distribution at higher alignment scores. The bottom panel illustrates the relationship between E-values (on a logarithmic scale, represented as 1e-X) and the proportion of positives to alignment length, showing a central peak around the midrange values, indicating optimal alignments with significant statistical relevance. This figure provides insights into sequence alignment characteristics, emphasizing patterns in sequence abundance, alignment quality, and statistical significance.

The distribution of E-values for sleep-related genes aids in predicting their roles and evolutionary trajectories. In bioinformatics, E-values are relative indicators of the alignment score of sequences and the statistical significance linked to the alignment of a sequence with a database of sequences. The distribution displays tendencies indicative of potential conserved domains, suggesting that these genes are crucial for regulating sleep and other biological processes (refer to Figure 1, Middle panel). Consequently, highly conserved sequences with low E-values are likely essential orthologs, evolutionarily preserved owing to their indispensable roles. In contrast, sequences with high E-values may be interpreted as adaptive modifications or species-specific components. These patterns aid in deducing the ecological, behavioral, or metabolic influences on sleep-related genes [27]. These results are essential for guiding functional investigations to investigate gene roles, particularly emphasizing conserved regions due to their likely significance in sleep. They also point to other sequences with limited homology that may influence organismal variation.

Figure 1 (bottom panel) provides data on the sequence similarity of genes associated with sleep processes. The graphical representation illustrates the distribution of sequence similarity levels, quantified as the ratio of positive hits to alignment length. The figure highlights frequency and emphasizes variances and irregular values at various thresholds. This information may assist in clarifying the genetic basis of sleep since it is feasible to identify the more conserved regions or those exhibiting significant sequence similarity within these genes.

### EGGNOG Annotation

Figure 2 (top panel) depicts the distribution of sequences (Seqs) using a bar chart representing five GO keywords labeled 0, 1, 2, 3, 4, and 5. This distribution reveals that most sequences in the examined set are closely associated with GO term 0, representing a significant biological category or process within the dataset. Others represent little or need more substantial association or relevance to the analyzed sequences. The differences may be important enough to illuminate the particular region of interest within the biological environment or data collection, such as a predominant cellular activity or an extensively examined or abundantly present structure.

**Figure 2.**
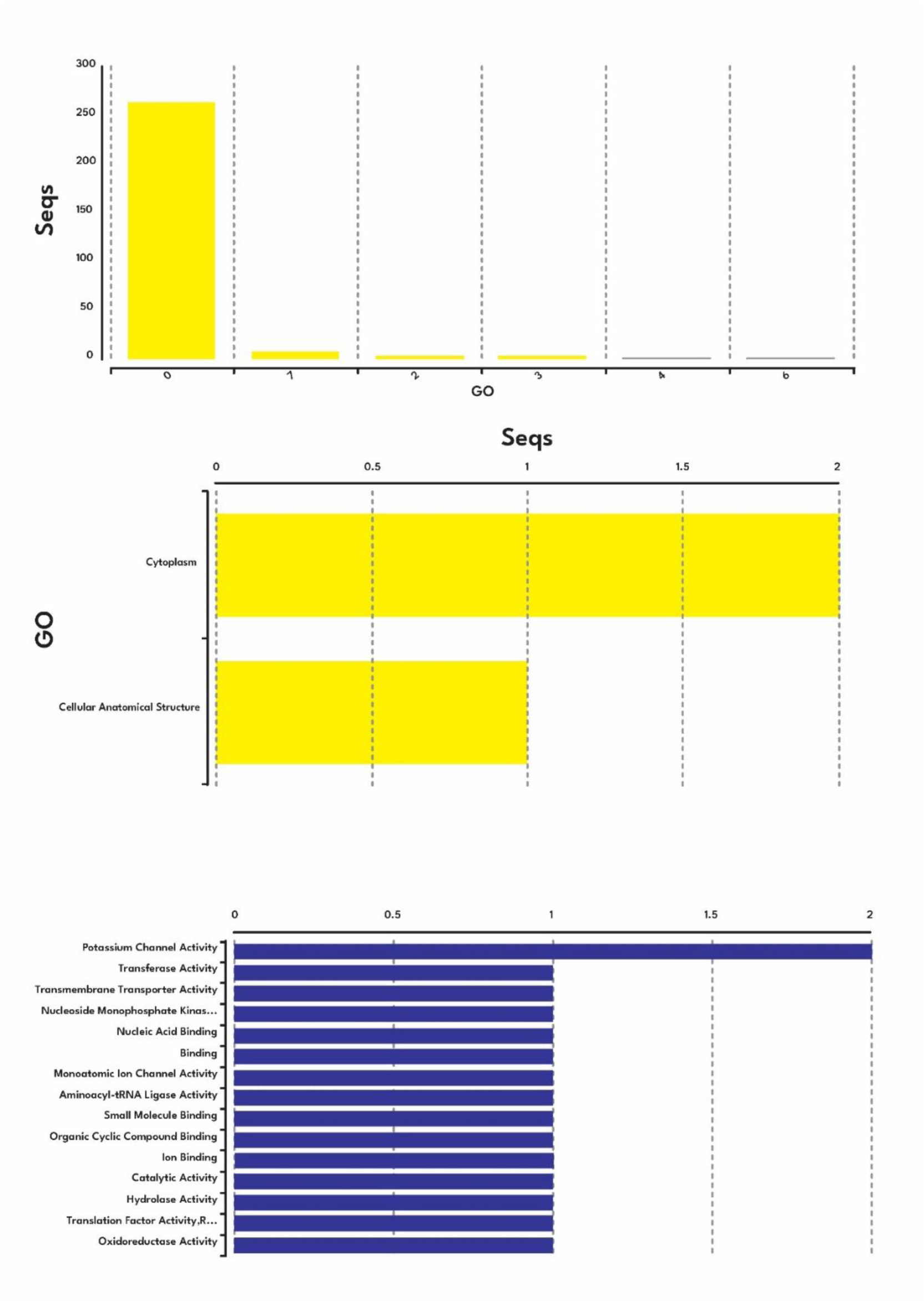
Sequence analysis using Gene Ontology (GO). The first data panel represents the number of sequences (Seqs) associated with a particular GO terms and the major bar at GO term 0 suggests the most dominant category. The middle panel concerns cellular localization and sequences closely linked to the cytoplasm and the cellular anatomical structures. The lowest panel shows molecular functions overrepresented in the dataset, which include potassium channel activity as the most representative function, followed by the transferase activity, nucleic acid binding, and ion binding functions. These findings give functional meanings and cellular localization to the sequences that have been identified.

Figure 2 (middle panel) depicts the allocation of sequences (Seqs) among two Gene Ontology (GO) terms: “cytoplasm” and “cellular anatomical structure.” The “cytoplasm” category comprises two sequences, which is quite near the total number of studied sequences, indicating a significant correlation between many sequences and this word. Conversely, the phrase “cellular anatomical structure” is less prevalent and approaches a frequency of one, indicating that such sequences are less represented in the collection of compared sequences. The distribution of this mode means that the biological components or processes encoded by the sequences are substantially linked to the cytoplasm and that the function/cell localization analysis used in the research may be pertinent to this feature. The limited number of papers in the “cellular, anatomical structure” category may result from a more stringent definition of cellular components or the term’s relatively low relevance within the overall collection. Further information on this matter would rely on the kind of data, treatment circumstances, research objectives, and the distribution curve of other Gene Ontology words to aid in comprehending the biological context.

Figure 2 (bottom panel) illustrates the relative abundance of sequences (Seqs) according to the gene ontology (GO), primarily focusing on Molecular Function. The most significant number of GO keywords is identified in the sequences related to ‘Potassium channel activity.’ Additional GO keywords such as “transferase activity,” “transmembrane transporter activity,” “nucleoside monophosphate kinase activity,” and various forms of “binding” and enzymatic activities exhibit comparable albeit somewhat reduced sequence counts. This distribution indicates that the predominant functional category in this dataset is ‘potassium channel activity,’ perhaps due to the prevalence of proteins associated with ion channels or their activities. The remaining GO keywords demonstrated adequate coverage of the dataset’s molecular functions, including various activities such as enzymatic roles, transferases, hydrolases, oxidoreductases, and molecular binding involving nucleic acids, ions, and small molecules. The presence of these activities in most of the investigated sequences, which are mostly identical with one exception, indicates the functional heterogeneity of the examined sequences. A further investigation may examine the biological environment, such as the processes or pathways involved in these functions.

### GO Level Distribution and Enzyme Classes

Figure 3 (top panel) depicts the sequence counts designated as ‘Seqs’ across four enzyme EC classes: Oxidoreductases, Transferases, Hydrolases, and Ligases. Transferases (EC 2) have the highest sequences, surpassing other enzyme classes. This indicates their importance in the dataset, likely owing to their role in the catalysis of functional groups such as methyl, glycosyl, or phosphate, which are essential for metabolism and signaling. This contrasts with Oxidoreductases, Hydrolases, and Ligases, which have comparable but quantitatively inferior absolute sequence counts. Hydrolases are crucial for catalyzing reactions that include the cleavage of a chemical bond by water, whereas oxidoreductases facilitate reactions involving the oxidation and reduction of molecules via redox processes. The numbers on the x-axis range from 0 to 2, representing the normalized or reduced sequence count of all classes. The prevalence of Transferases indicates that the detected proteins may have a functional or biological role relevant to the dataset; hence, other systems, pathways, or environmental circumstances associated with the dataset warrant further exploration.

**Figure 3.**
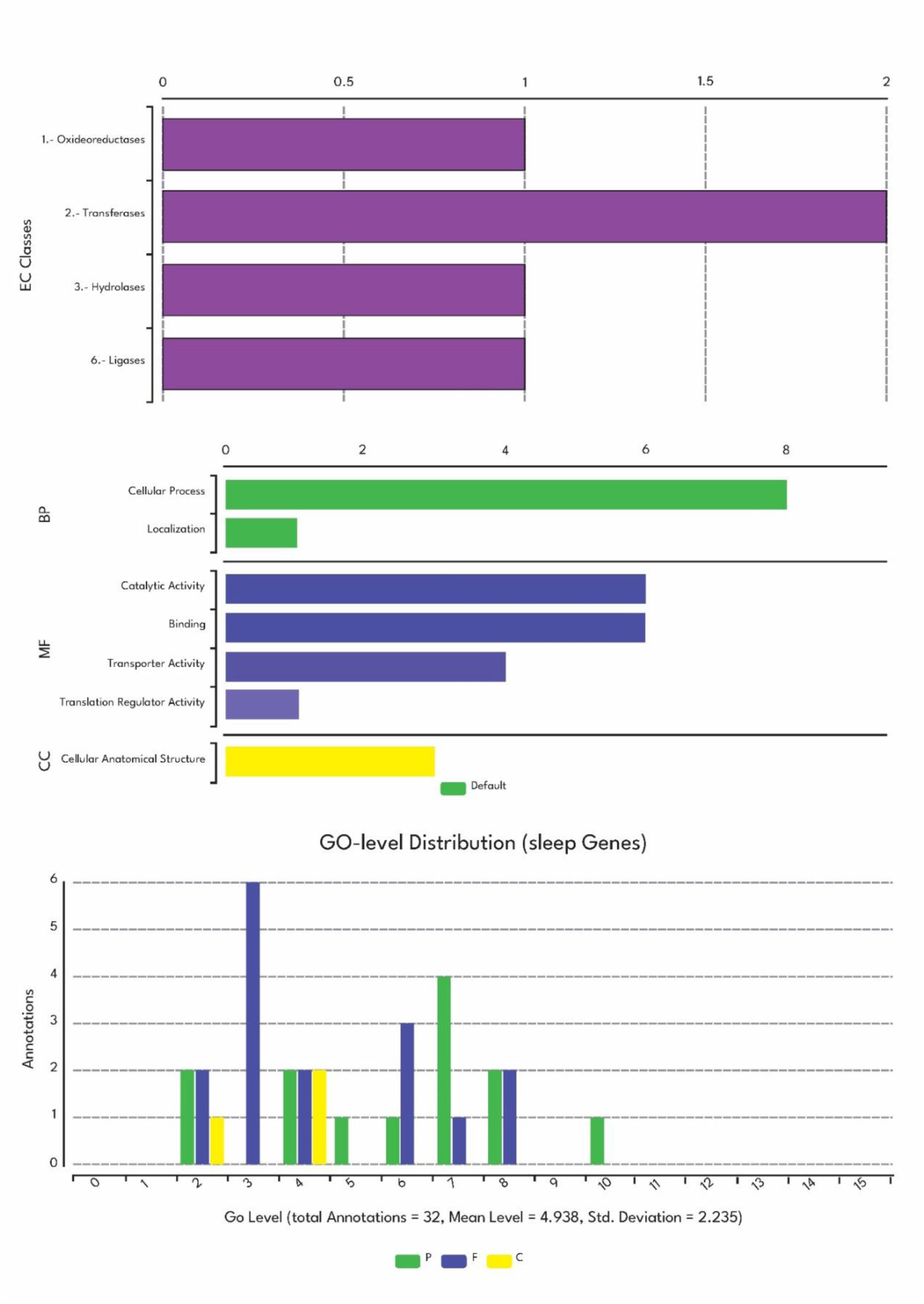
Functional classification and Gene Ontology level distribution of sleep-related genes. The enzyme classes (EC Classes) read in the top panel show that we have transferases, hydrolases, ligases, and oxidoreductases among others. The middle panel shows GO annotations where the data is classified under BP, MF, and CC. At the level of the biological processes, the cellular processes are the most represented while the molecular functions including the catalytic activity and binding show the highest enrichment. In the cellular component category, it is also observed that the structures of cell anatomy dominate. The bottom panel shows the distribution of the sleep-related genes at the GO level showing the count of annotations in various GO levels. The distribution shows the functional and structural diversity of genes related to sleep with a total of 32 annotations, an average of 4.938 ± 2.235.

Figure 3 (middle panel) depicts a horizontal bar chart that summarizes the distribution of sequences (referred to as “Seqs”) across three primary Gene Ontology (GO) categories: Biological Process, Molecular Function, and Cellular Component. In the first subcategory, Biological Process (BP), the phrase “cellular process” occurs four times more often than “localization.” This suggests that most examined sequences participate in fundamental cellular activities, such as metabolic processes, signaling, or cell division. Still, a few are associated with localization processes that include the movement of molecules inside the cell. The predominant GO keywords identified in the Molecular Function (MF) category are ‘catalytic activity’ and ‘binding,’ followed by ‘transporter activity’ and ‘translation regulator activity.’ The predominance of catalytic activity and binding indicates that most sequences are enzymes or molecules capable of binding substrates or ligands. Transporter activity is evident, suggesting roles in the translocation of molecules across membranes, whereas translation regulator activity, albeit less prevalent, indicates involvement in the regulation of translation.

All sequences in the Cellular Component (CC) category pertain to ‘cellular anatomical structures,’ indicating the cellular compartments or structures where these proteins are localized, as shown in Figure 3 (bottom panel). This extensive category aligns with their functional and biological significance. The graphic effectively illustrates the functional diversity of the examined sequences, their role in fundamental cellular processes and interactions with other molecules, and the structural positioning system. This study might be enhanced by using more specific GO keywords to elucidate these sequences’ more refined functional and biological settings.

### INTERPRO Analysis

Figure 4 (top panel) presents a bar graph illustrating the distribution of protein sequences across various IPS domains as analyzed by InterProScan. The proportion of sequences in the “others” category is much more significant than the other categories. This suggests several sequences are associated with low-frequency IPS domains or domains not included in the table. Among the enumerated domains, the GPCR, rhodopsin-like (IPR017452) domain had the most significant sequence count, suggesting it may be the most frequent or essential domain within the dataset. Other significant domains include the Ion transport domain, Pfam number PF014387, and the Myc-type, basic helix-loop-helix domain, Pfam number PF03643; the majority of the other mentioned domains have a relatively low proportion of sequence occurrence. This distribution illustrates the outliers among the data points, mainly concentrated within a few domains, with a category labeled ‘others’ that requires more investigation to ascertain its composition.

**Figure 4.**
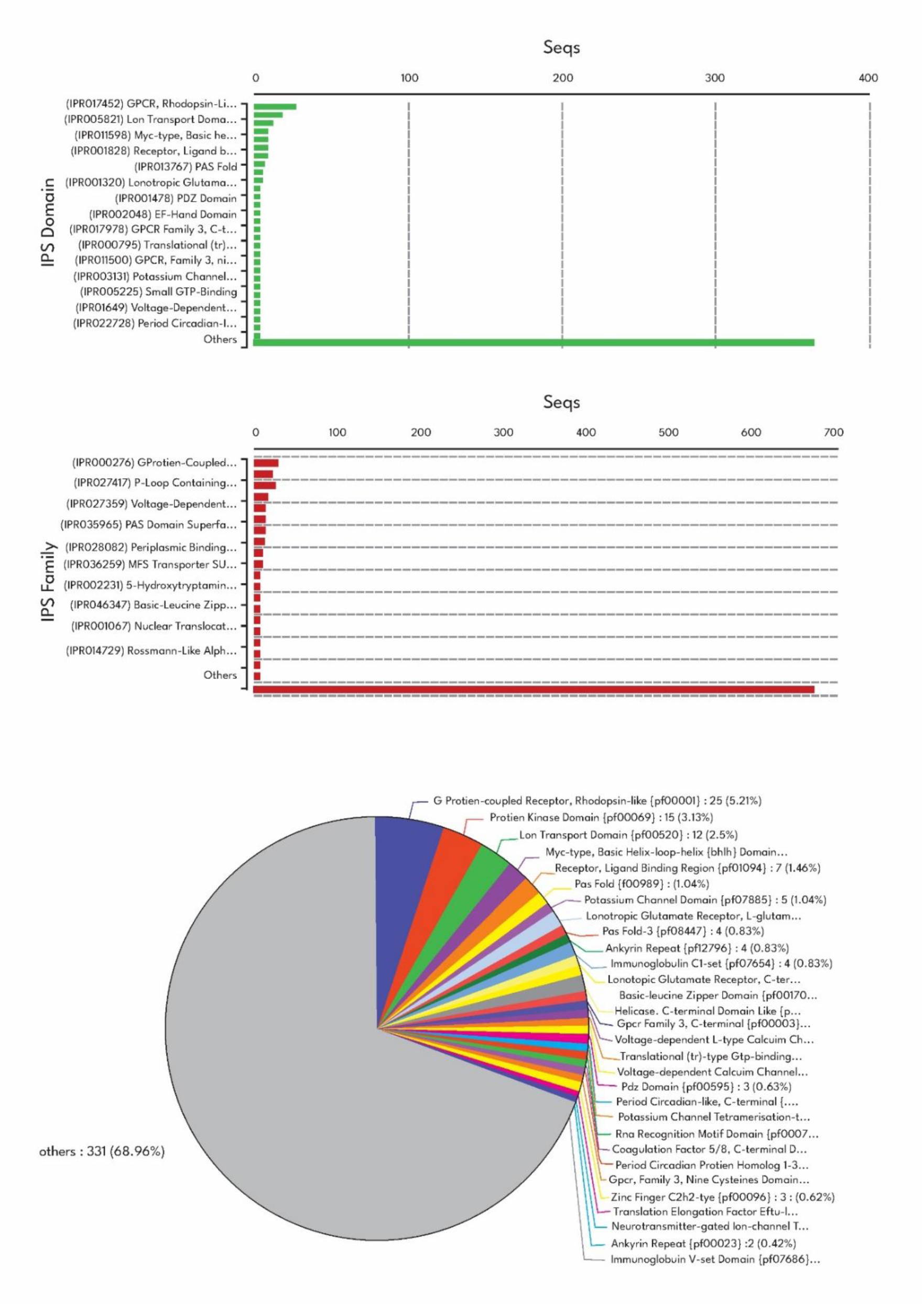
The distribution of the protein sequences concerning IPS domains and families. The first graph shown in the figure represents the abundance of sequence in various IPS domains where the largest domain was the GPCR domain (IPR017452). The middle panel shows the distribution of the frequency of sequences identified to IPS families, and the G-protein coupled receptor family IPS000276 was found significantly overrepresented. The bottom panel shows a pie chart indicating the ratio of different protein families – some families are dominant and the ‘Other’ is as large as 68.96% of the sequences.

Figure 4 (middle panel) depicts a bar graph representing the number of protein sequences (Seqs) and the number of InterProScan (IPS) families. The “others” group is the most significant portion of the data, indicating that several sequences are linked to infrequent IPS families or unrelated to specifically listed families. Among the particular IPS families, IPR000276, representing the G protein-coupled receptors superfamily, has the highest number of sequences, reinforcing its dominance or pervasiveness throughout the dataset. The further families include P-loop, including nucleoside triphosphate hydrolase (PS51180) and voltage-dependent calcium channel (PS50806), whereas the remaining families have little representation. This distribution emphasizes the substantial number of families in the data set and the considerable quantity of unidentified families, sometimes referred to as ‘others.’

Figure 4 (bottom panel) illustrates the Pfam protein families associated with the sequences examined in the research. The predominant segment of the identified sequences is categorized as “others” - including 331 sequences, or 68.96% of the overall total; this indicates the presence of several sequences from families that are either uncommon or not shown on the chart. The most prominent family identified is the G protein-coupled receptor, rhodopsin-like (PF00001), with 25 sequences (5.21%), indicating its significant presence or functional importance within the dataset. Additional families include the Protein kinase domain, with 15 sequences (3.13%) for PF00069, and the Ion transport domain, with 12 sequences (2.5%) for PF00520. Most families include a minimal fraction of the overall sequences, equivalent to or less than 2%, including the Myc-type essential helix-loop-helix domain (PF00010) and the PAS fold domain (PF00989). The distribution shown here indicates that some protein families are well-represented, although a significant majority of the sequences are spread throughout several comparably underrepresented protein families. The “others” category contains additional sequences that suggest a greater degree of sequence divergence, necessitating investigation to identify potentially overrepresented or previously unrecognized protein families.

### Pathway Analysis

Figure 5 depicts the allocation of pathways among six KEGG (Kyoto Encyclopedia of Genes and Genomes) categories: Somatic diseases, Organism systems, Enterprise organisms, Adaptation, Defense, Communication, Transportation, Membrane transport, Cell growth and function, and Signal transduction. The “Found” prefixes are the most prevalent across all categories, with Human Diseases and Organismal Systems being the two most represented, each including roughly 90 routes. Nevertheless, the quantity of “Enriched” routes across all categories is minimal, indicating a lack of statistical significance in the enrichment study. Metabolic pathways have a substantial participation score, including over 60 pathways, followed by Environmental Information Processing with around 30 pathways and Cellular Processes with about 20 pathways. Genetic Information Processing contains the fewest routes, around ten, indicating that this function contributes little to the dataset. The Human Diseases and Organismal Systems pathways are predominant, signifying the dataset’s relevance in clinical and physiological contexts, with a substantial representation of metabolic pathways crucial for biological activities. The lack of significant overrepresentation for any category may result from a very diversified dataset with an equitable distribution of paths or an insufficient statistical overrepresentation value. This distribution underscores the biological significance of the dataset, indicating that more efforts may be required to refine the enrichment levels or to analyze the patterns of specific pathways. This figure demonstrates the potential of the dataset for exploring other aspects of the biomedical domain and many facets of biology and disorders.

**Figure 5.**
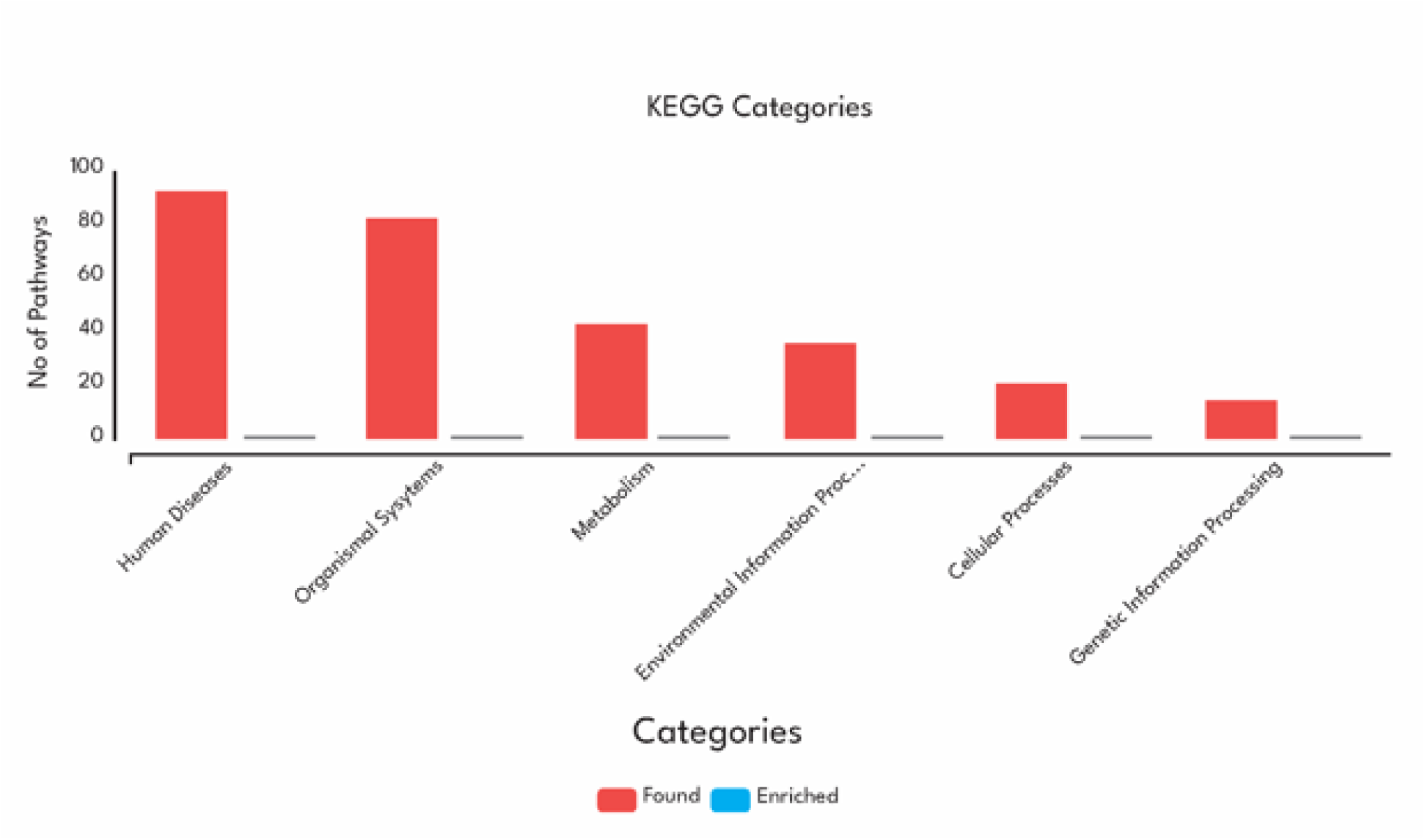
Distribution of pathways across KEGG categories. The figure shows the number of pathways identified (“Found”) and statistically significant pathways (“Enriched”) across six KEGG categories: Human Diseases, Organismal Systems, Metabolism, Environmental Information Processing, Cellular Processes, and Genetic Information Processing. The “Found” pathways are most abundant in the Human Diseases and Organismal Systems categories, while enrichment is minimal across all categories. This distribution highlights the dataset’s focus on disease-related and organism-level processes, with a broad representation of metabolic pathways and limited enrichment significance.

## DISCUSSION

This study provides a unique phylogenomic methodology for elucidating the evolutionary links and functional implications of sleep-related genes. Our findings enhance comprehension of sleep-related processes’ evolutionary and functional adaptability, particularly in organisms within the Epsilonproteobacteria Class [28]. We compared and contrasted our findings to previous phylogenomic investigations. The finding identifies sleep orthologs in *S. paralvinellae*, including the molecular chaperone DnaK, serine hydroxymethyltransferase, and potassium channel family proteins. These genes exhibited a significant degree of sequence similarity recognized for their involvement in sleep regulation across several species, with mean values ranging from 39.13% to 61.45%. This aligns with previous research on sleep orthologs, which found significant conservation of sleep-associated genes, including chaperones and ion channels. The preservation of DnaK indicates its functions in protein folding and cellular repair, which are crucial during sleep and have previously been associated with stress response pathways.

In addition to these important findings, the study also discovered that small proteins, such as adenylate kinase, exhibited more significant conservation in their functional domains than multi-domain lengthy proteins [29]. This pattern corroborates earlier results that proteins with fewer domains exhibit greater conservation, suggesting they have undergone significant selective pressure due to their essential roles. The recurrent identification of bifunctional protein-serine/threonine kinases and phosphatases, suggests that *S. paralvinellae* possesses a unique adaptation capability potentially linked to its extremophile habitat. This finding suggests that these enzymes may play a role in assisting adaptation to environmental conditions, inasmuch as similar multifunctional proteins have been documented to participate in metabolism under harsh settings. Our Gene Ontology (GO) analysis identified cellular processes, catalytic activities, and potassium (K+) channel activities as the most enriched functional categories among the sleep-related genes. The augmented K+ channel activity aligns with prior research showing that ion channels play a role in sleep-wake transitions [30–33] The finding of sequences linked to transmembrane transporter function and nucleoside monophosphate kinase activity suggests that additional pathways exist through which these genes may participate in sleep-related processes, including energy metabolism and molecular transport.

The pathway analysis unexpectedly revealed that high enrichment occurred in metabolic and signal transduction pathways, whereas enrichment happened less frequently in sleep-related pathways. This is consistent with the recognition that numerous sleep-related genes possess universal metabolic functions, a characteristic noted in other model species such as *D. melanogaster* and *C. elegans* [34,35]. Nonetheless, the inability to identify statistically significant pathways may indicate that the metabolisms and ecological niche of *S. paralvinellae* diverge from current understandings of sleep regulation. Our study further underscores the complexity of orthologous relationships. While specific sleep-related genes appear to retain conserved functions across species, others demonstrate signs of subfunctionalization [36] (i.e., the division of various subsets of an ancestor gene’s functions among daughter copies during gene duplication is known as subfunctionalization. For example, duplicate genes will share redundant roles that overlap early in the process of functional divergence, without each duplication event having to confer a selection advantage.) or neofunctionalization [37]. For instance, bifunctional protein-serine/threonine kinases identified in *S. paralvinellae* may represent an adaptive divergence, reflecting its evolutionary history within the extreme conditions of hydrothermal vents. This observation complements earlier findings that orthologs can evolve distinct functions over time, particularly in response to environmental pressures.

The functional divergence of orthologs is a critical consideration for understanding sleep mechanisms. As highlighted by Gabaldón and Koonin (2013) [38], orthologous genes often exhibit nuanced differences in regulatory or structural domains that allow for species-specific adaptations. Our findings suggest that, while core sleep-related processes are conserved, the specific regulatory mechanisms may vary significantly depending on ecological and evolutionary context. Ultimately, our research substantiates the notion that orthologous relationships are complex. Specific sleep-related genes appear to have preserved their ancestral functions, although others exhibit signs of subfunctionalization or neofunctionalization. Bifunctional protein-serine/threonine kinases in *S. paralvinellae* may represent an adaptive divergence resulting from its evolution in the stressful environment of hydrothermal vents. These data corroborate other studies indicating that orthologs can evolve diverse functions throughout time in response to environmental pressures [39,40].

The divergence of orthologs is a crucial element of functional differentiation in sleep research. Gabaldón and Koonin (2013) posited that orthologous genes may exhibit modest alterations in their regulatory or structural areas, facilitating species-specific variants [38]. Our data suggest that fundamental sleep-regulating mechanisms are analogous across species. However, the regulatory systems may vary in distinct ecological and evolutionary contexts. This study illustrates the utility of phylogenomic analysis in elucidating the evolutionary history of sleep-associated genes. This study utilized extensive genomic data to delineate how conserved characteristics may be linked to specialized adaptations in *S. Paralvinellae*. These findings contribute to a more comprehensive understanding of the intricate connections among sleep, metabolism, and environmental adaption.

Future research should involve experimental confirmation of the identified genes and pathways in both model and non-model organisms. Moreover, the phylogenomic analysis might be expanded to additional Epsilonproteobacteria species to enhance understanding of sleep’s evolutionary history and functional adaptability. Comparisons with more thoroughly studied model species, such as *Hydra vulgaris* and Cassiopea, may aid in bridging the gap between phylogenomic hypotheses and the phenotypic manifestation of sleep-related activities [9].

Whether certain species sleep at all is still a controversial and frequently disputed hypothesis. There is insufficient empirical evidence to support the claim that schools of fish and pelagic cruisers do not require sleep [41–43]. The current investigation suggests that such controversies might be resolved through phylogenetic and phylogenomic analysis, which in turn could point to sleep’s evolutionary origin and function.

The study of phylogeny, or the evolutionary history of organisms, can aid in the understanding of how they have evolved throughout evolution and how they are related to one another. Within the discipline of evolutionary biology, phylogenetics and phylogenomics examine the links between organisms and the evolution of diseases. However, these disciplines differ in scope, methodology, and the types of data they utilize. Therefore, the goal of both phylogenetics and phylogenomics is to provide a more integrated view of these phenomena. They do this by reconstructing evolutionary links among species or genes; phylogenetic trees, which are the descriptive outcomes of this analysis, usually represent these relationships.

While determining evolutionary relationships from molecular data (such as DNA, RNA, or protein sequences) is the common goal of both phylogenetics and phylogenomics, phylogenomics offers a more thorough and genome-wide approach, utilizing large-scale datasets to provide more profound and more detailed insights into evolutionary history. Additionally, phylogenomics provides a broader understanding of the evolution of phenotypic features, behaviors, and diseases. It offers a potent framework for comprehending the genetic foundations of behaviors (e.g., genetic influence on behavior, gene regulation, and expression), diseases (e.g., pathogen evolution, host-pathogen interaction, the genetic basis of disease susceptibility), and phenotypic traits (e.g., gene-phenotype association, ancestral trait reconstruction, and identify the genetic basis of convergent evolution). Researchers may follow the evolutionary history of these phenotypic traits and gain insights into their emergence, diversification, and adaptation across time by utilizing large-scale genomic data. Therefore, such an approach is extremely helpful in resolving complex questions in evolutionary biology, medicine, and other relevant sciences.

The claim that an ortholog will only serve one function is also untrue [38]. Genes in distinct species that diverged from a common ancestor through speciation are known as orthologs. These orthologs can diverge functionally over time, even if they frequently maintain the same function. Under this process, which is referred to as “subfunctionalization” or “neofunctionalization,” orthologous genes may take on new or different activities [40].

Therefore, orthologous genes are not limited to a single function, and orthologs can diverge in function across evolutionary time scales. Such knowledge is imperative for interpreting gene function across species since different creatures may use the same gene to support distinct biological processes.

It should be noted that the study has certain limitations. First, a lab-based confirmation is needed for the computational predictions. Second, an experimental confirmation of identified genes and pathways is required. Furthermore, the study results cannot be generalizable as the study findings may or may not applicable to non-extremophilic species.

Despite these limitations, this study offers unique insights beyond the conventional animal models used in sleep research because it is the first to examine sleep-related genes in extremophilic bacteria. For a thorough gene analysis, the study used phylogenomics tools and methods like BLAST, GO annotations, and KEGG pathways. According to the study, conserved genes (such as potassium channels and DnaK) are crucial in regulating sleep. Furthermore, bifunctional enzymes’ functional divergence indicates environmental adaptation. Most significantly, the study links the fields of sleep research of multicellular organisms with bacterial genomics.

## CONCLUSION

This work establishes a solid framework for investigating the evolutionary history and functional importance of sleep genes across diverse taxa. On this note, this study illustrates the significance of phylogenomic investigations in clarifying the evolutionary and functional characteristics of sleep genes of *S. paralvinellae* of Epsilonproteobacteria Class. This study demonstrates that essential sleep genes, such as DnaK and potassium channel family proteins, are highly conserved across evolutionary history. The elevated conservation observed in tiny, single-domain proteins such as adenylate kinase suggests that their functions are significant under specific selective pressures. Moreover, the finding of protein-serine/threonine kinases and phosphatases reinforces the concept of functional diversification in *S. paralvinellae* within its extremophile environment, influenced by ecological processes. The enrichment analysis revealed that the primary functions connected with metabolism and signal transduction indicated the involvement of sleep genes in overall metabolism. Nonetheless, shared basic processes are apparent in sleep regulation, yet subfunctionalization and neofunctionalization in certain orthologs indicate species-specific variations. These findings expand the understanding of sleep evolution and its connection to metabolic and environmental adaptability, illustrating the interplay between inherent genes and ecological influences. Subsequent research should encompass experimental validation of these results in both model and non-model organisms, along with comparative analyses of other members of the Epsilonproteobacteria group, to enhance comprehension of the evolutionary and functional intricacies of sleep-related processes.

## CRediT authorship contribution statement

SRP; SBC: Conceptualization and Methodology; SRP; KMS; SP: Data collection and review, KMS; SP: Software, SRP; KMS; SP: Preparation of original draft; DWS; SBC: Critical review and editing; SBC: supervision; Prior to submission, all authors (SRP; KMS; SP; DWS; SBC) have read and agreed to the submitted version of the manuscript.

## Author Agreement Statement

All authors explicitly confirm that the paper is their original work that has not previously been published nor is it presently being considered for publication in any other journal, venue, or context. Each author expressly declares that they contributed equally, read, evaluated, and accepted the final version of the paper, and agreed to be included as co-authors per ICMJE guidelines. Each of the authors has approved the author sequence and the corresponding authors.

## Role of medical writer or editor

Not applicable.

## Ethical Statement

This study does not contain any work involving animals or human participants performed by any of the authors. Hence, no IRB approval was necessary for this work.

## Data availability

This study utilized the dataset originally prepared for our ongoing phylogenomics and bioinformatics investigation of sleep. Since this is a work in progress, the authors cannot provide additional information.

## References

1. Pandi-Perumal SR, Saravanan KM, Paul S, et al. (2024) Waking Up the Sleep Field: An Overview on the Implications of Genetics and Bioinformatics of Sleep. Mol Biotechnol 66: 919–931.

2. Pandi-Perumal SR (2010) Great challenges to sleep medicine: problems and paradigms. Front Neurol 1: 7.

3. Libourel P-A, Herrel A (2016) Sleep in amphibians and reptiles: a review and a preliminary analysis of evolutionary patterns. Biol Rev 91: 833–866.

4. Campbell SS, Tobler I (1984) Animal sleep: A review of sleep duration across phylogeny. Neurosci Biobehav Rev 8: 269–300.

5. McNamara P, Barton RA, Nunn CL (Eds.) (2010) Evolution of sleep: Phylogenetic and functional perspectives., New York, NY, US, Cambridge University Press.

6. Karmanova IG (1982) Evolution of Sleep: Stages of the Formation of the ‘Wakefulness-Sleep’ Cycle in Vertebrates Translation from Russian by A.I. Koryushkin and O.P. Uhastkin, Leningrad With Editorial Assistance by P. Koella (Basel).

7. Hendricks JC, Sehgal A, Pack AI (2000) The need for a simple animal model to understand sleep. Prog Neurobiol 61: 339–351.

8. Pandi-Perumal SR, Saravanan KM, Paul S, et al. (2024) Harnessing Simple Animal Models to Decode Sleep Mysteries. Mol Biotechnol.

9. Kanaya HJ, Park S, Kim J, et al. (2024) A sleep-like state in Hydra unravels conserved sleep mechanisms during the evolutionary development of the central nervous system. Sci Adv 6: eabb9415.

10. Nath RD, Bedbrook CN, Abrams MJ, et al. (2017) The Jellyfish *Cassiopea* Exhibits a Sleep-like State. Curr Biol 27: 2984–2990.e3.

11. Beeby M (2015) Motility in the epsilon-proteobacteria. Curr Opin Microbiol 28: 115–121.

12. Jaeschke A, Jørgensen SL, Bernasconi SM, et al. (2012) Microbial diversity of Loki’s Castle black smokers at the Arctic Mid-Ocean Ridge. Geobiology 10: 548–561.

13. Takai K, Suzuki M, Nakagawa S, et al. (2006) Sulfurimonas paralvinellae sp. nov., a novel mesophilic, hydrogen- and sulfur-oxidizing chemolithoautotroph within the Epsilonproteobacteria isolated from a deep-sea hydrothermal vent polychaete nest, reclassification of Thiomicrospira denitrificans as Su. Int J Syst Evol Microbiol 56: 1725–1733.

14. Motschall E, Falck-Ytter Y (2005) Searching the MEDLINE Literature Database through PubMed: A Short Guide. Onkologie 28: 517–522.

15. Barshir R, Fishilevich S, Iny-Stein T, et al. (2021) GeneCaRNA: A Comprehensive Gene-centric Database of Human Non-coding RNAs in the GeneCards Suite. J Mol Biol 433: 166913.

16. Hamosh A, Amberger JS, Bocchini C, et al. (2021) Online Mendelian Inheritance in Man (OMIM®): Victor McKusick’s magnum opus. Am J Med Genet Part A 185: 3259–3265.

17. Pandi-Perumal SR, Saravanan KM, Paul S, et al. (2025) Unraveling the Mysteries of Sleep: Exploring Phylogenomic Sleep Signals in the Recently Characterized Archaeal Phylum Lokiarchaeota near Loki’s Castle. Int J Mol Sci 26.

18. Altschul SF, Madden TL, Schäffer AA, et al. (1997) Gapped BLAST and PSI-BLAST: a new generation of protein database search programs. Nucleic Acids Res 25: 3389–3402.

19. Huerta-Cepas J, Forslund K, Coelho LP, et al. (2017) Fast Genome-Wide Functional Annotation through Orthology Assignment by eggNOG-Mapper. Mol Biol Evol 34: 2115–2122.

20. Kanehisa M, Goto S (2000) KEGG: Kyoto Encyclopedia of Genes and Genomes. Nucleic Acids Res 28: 27–30.

21. Makarova KS, Sorokin A V, Novichkov PS, et al. (2007) Clusters of orthologous genes for 41 archaeal genomes and implications for evolutionary genomics of archaea. Biol Direct 2: 33.

22. Blum M, Chang H-Y, Chuguransky S, et al. (2021) The InterPro protein families and domains database: 20 years on. Nucleic Acids Res 49: D344–D354.

23. Milacic M, Beavers D, Conley P, et al. (2024) The Reactome Pathway Knowledgebase 2024. Nucleic Acids Res 52: D672–D678.

24. Chojnowski G, Bochtler M (2007) The statistics of the highest {\it E} value. Acta Crystallogr Sect A 63: 297–305.

25. Fourie KR, Wilson HL (2020) Understanding GroEL and DnaK Stress Response Proteins as Antigens for Bacterial Diseases. Vaccines 8.

26. Gallego M, Virshup DM (2005) Protein serine/threonine phosphatases: life, death, and sleeping. Curr Opin Cell Biol 17: 197–202.

27. van Rosmalen L, Deota S, Maier G, et al. (2024) Energy balance drives diurnal and nocturnal brain transcriptome rhythms. Cell Rep 43: 113951.

28. Waite DW, Vanwonterghem I, Rinke C, et al. (2017) Comparative Genomic Analysis of the Class Epsilonproteobacteria and Proposed Reclassification to Epsilonbacteraeota (phyl. nov.). Front Microbiol 8.

29. Solaroli N, Panayiotou C, Johansson M, et al. (2009) Identification of two active functional domains of human adenylate kinase 5. FEBS Lett 583: 2872–2876.

30. Yoshida K, Shi S, Ukai-Tadenuma M, et al. (2018) Leak potassium channels regulate sleep duration. Proc Natl Acad Sci 115: E9459–E9468.

31. Kodirov S (2022) Functioning of K channels during sleep. Arch Insect Biochem Physiol 110.

32. Steinberg EA, Wafford KA, Brickley SG, et al. (2015) The role of K2P channels in anaesthesia and sleep. Pflügers Arch - Eur J Physiol 467: 907–916.

33. Pang DSJ, Robledo CJ, Carr DR, et al. (2009) An unexpected role for TASK-3 potassium channels in network oscillations with implications for sleep mechanisms and anesthetic action. Proc Natl Acad Sci 106: 17546–17551.

34. Harbison ST, McCoy LJ, Mackay TFC (2013) Genome-wide association study of sleep in Drosophila melanogaster. BMC Genomics 14: 281.

35. Bringmann H (2019) Genetic sleep deprivation: using sleep mutants to study sleep functions. EMBO Rep 20: e46807.

36. Cusack BP, Wolfe KH (2007) When gene marriages don’t work out: divorce by subfunctionalization. Trends Genet 23: 270–272.

37. Kuzmin E, Taylor JS, Boone C (2022) Retention of duplicated genes in evolution. Trends Genet 38: 59–72.

38. Gabaldón T, Koonin E V (2013) Functional and evolutionary implications of gene orthology. Nat Rev Genet 14: 360–366.

39. Martijn J, Vosseberg J, Guy L, et al. (2018) Deep mitochondrial origin outside the sampled alphaproteobacteria. Nature 557: 101–105.

40. Zhang J (2003) Evolution by gene duplication: an update. Trends Ecol Evol 18: 292–298.

41. Kavanau JL (2008) Sleepless in the Sea. Science (80- ) 322: 527.

42. Lee Kavanau J (2005) Evolutionary approaches to understanding sleep. Sleep Med Rev 9: 141–152.

43. Kavanau JL (1997) Origin and Evolution of Sleep: Roles of Vision and Endothermy. Brain Res Bull 42: 245–264.

